# A Critical Role for Perivascular Cells in Amplifying Viral Haemorrhage Induced by Dengue Virus Non-Structural Protein 1

**DOI:** 10.1101/2020.02.13.948208

**Authors:** Yin P. Cheung, Valeria Mastrullo, Davide Maselli, Teemapron Butsabong, Paolo Madeddu, Kevin Maringer, Paola Campagnolo

## Abstract

Dengue is the most prevalent arthropod-borne viral disease affecting humans, with severe dengue typified by potentially fatal microvascular leakage and hypovolaemic shock. Blood vessels of the microvasculature are composed of a tubular structure of endothelial cells ensheathed by perivascular cells (pericytes). Pericytes support endothelial cell barrier formation and maintenance through paracrine and contact-mediated signalling, and are critical to microvascular integrity. Pericyte dysfunction has been linked to vascular leakage in noncommunicable pathologies such as diabetic retinopathy, but has never been linked to infection-related vascular leakage. Dengue vascular leakage has been shown to result in part from the direct action of the secreted dengue virus (DENV) non-structural protein NS1 on endothelial cells. Using primary human vascular cells, we show here that NS1 also causes pericyte dysfunction, and that NS1-induced endothelial hyperpermeability is more pronounced in the presence of pericytes. Notably, NS1 specifically disrupted the ability of pericytes to support endothelial cell function in a 3D microvascular assay, with no effect on pericyte viability or physiology. These effects are mediated at least in part through contact-independent paracrine signals involved in endothelial barrier maintenance by pericytes. We therefore identify a role for pericytes in amplifying NS1-induced microvascular hyperpermeability in severe dengue, and thus show that pericytes can play a critical role in the aetiology of an infectious vascular leakage syndrome. These findings open new avenues of research for the development of drugs and diagnostic assays for combating infection-induced vascular leakage, such as severe dengue.

**SIGNIFICANCE STATEMENT:** The World Health Organisation considers dengue one of the top ten global public health problems. There is no specific antiviral therapy to treat dengue virus and no way of predicting which patients will develop potentially fatal severe dengue, typified by vascular leakage and circulatory shock. We show here that perivascular cells (pericytes) amplify the vascular leakage-inducing effects of the dengue viral protein NS1 through contact-independent signalling to endothelial cells. While pericytes are known to contribute to noncommunicable vascular leakage, this is the first time these cells have been implicated in the vascular effects of an infectious disease. Our findings could pave the way for new therapies and diagnostics to combat dengue, and potentially other infectious vascular leakage syndromes.

## INTRODUCTION

Dengue virus (DENV) is a serocomplex of four viruses (DENV-1-4) and the most prevalent arthropod-borne virus (arbovirus) affecting humans. Almost half the world’s population lives in at-risk areas across the tropics and subtropics, with an estimated 390 million infections and 96 million symptomatic cases per year (1). Dengue patients experience a range of symptoms including high fever, leucopoenia, maculopapular rash, retro-orbital pain, arthralgia and myalgia (2, 3). A minority of patients (approximately 500,000 cases per year) develop severe dengue, typified by cardiovascular complications such as plasma leakage (e.g. pleural effusion, oedema) and/or bleeding (haemorrhage) manifesting during defervescence that can lead to hypovolaemic shock, organ failure and death (2, 3). Fluid replacement therapy and other supportive treatments reduce the death rate, but there are no specific drugs to treat dengue and no diagnostic measure to predict the development of the life-threatening severe dengue (2-4).

At a microvascular level, the precise mechanisms leading to vascular leakage in severe dengue remain incompletely understood. A dysregulated cytokine response has been proposed to be a major contributor to vascular hyperpermeability, especially in heterotypic secondary infections with a different DENV serotype to the first infection (3, 5, 6). Antibodies against the viral non-structural protein NS1 have also been shown to bind to endothelial cells in secondary infections, causing their apoptosis (7-9). In addition, NS1 protein is abundantly secreted into patient serum and has been shown to directly induce endothelial cell hyperpermeability (10, 11). Serum NS1 concentrations in dengue patients vary widely. Although concentrations as high as 15,000 ng/ml have been recorded in some patients, concentrations of 10-1,000 ng/ml are more typical, with one study suggesting that levels above 600 ng/ml may be predictive of severe disease (12-14). NS1 enters endothelial cells through dynamin- and clathrin-dependent endocytosis and induces the expression of secreted cellular sialidases, heparanase and cathepsin L that cleave components of the endothelial cell glycocalyx reducing the integrity of the barrier (15-18). Inhibitors of sialidases and heparanases reduce NS1-dependent endothelial hyperpermeability *in vitro*, and NS1 vaccination has been shown to protect against vascular leakage *in vivo* (11, 17).

The published mechanistic studies into NS1-induced dengue vascular leakage primarily assessed endothelial cell function. However, the vessels affected in severe dengue *in vivo*, primarily capillaries and post-capillary venules, are comprised of an endothelial cell lining surrounded by perivascular cells (pericytes) embedded within the basement membrane (Fig 1A) (19, 20). There is a dynamic relationship between pericytes and endothelial cells, with two-way paracrine signalling as well as contact-mediated communication facilitated by pericyte pseudopod extensions that wrap around the endothelial cells (21). Pericytes are essential in both blood vessel formation to drive endothelial cell migration, proliferation and maturation, and in homeostasis for the maintenance and regulation of the endothelial barrier in established vessels (22, 23). Pericyte deficiency is perinatally lethal in mouse models due to widespread vascular leakage and aneurysms (24). In established vessels, pericyte dysfunction leads to microvascular hyperpermeability, such as in diabetic retinopathy where chronic exposure to heightened glucose levels due to unmanaged diabetes causes capillary occlusions, microaneurysms and blindness (25). To date, the role of pericytes in the aetiology of vascular hyperpermeability caused by an infectious disease has not been described.

**Fig 1.**
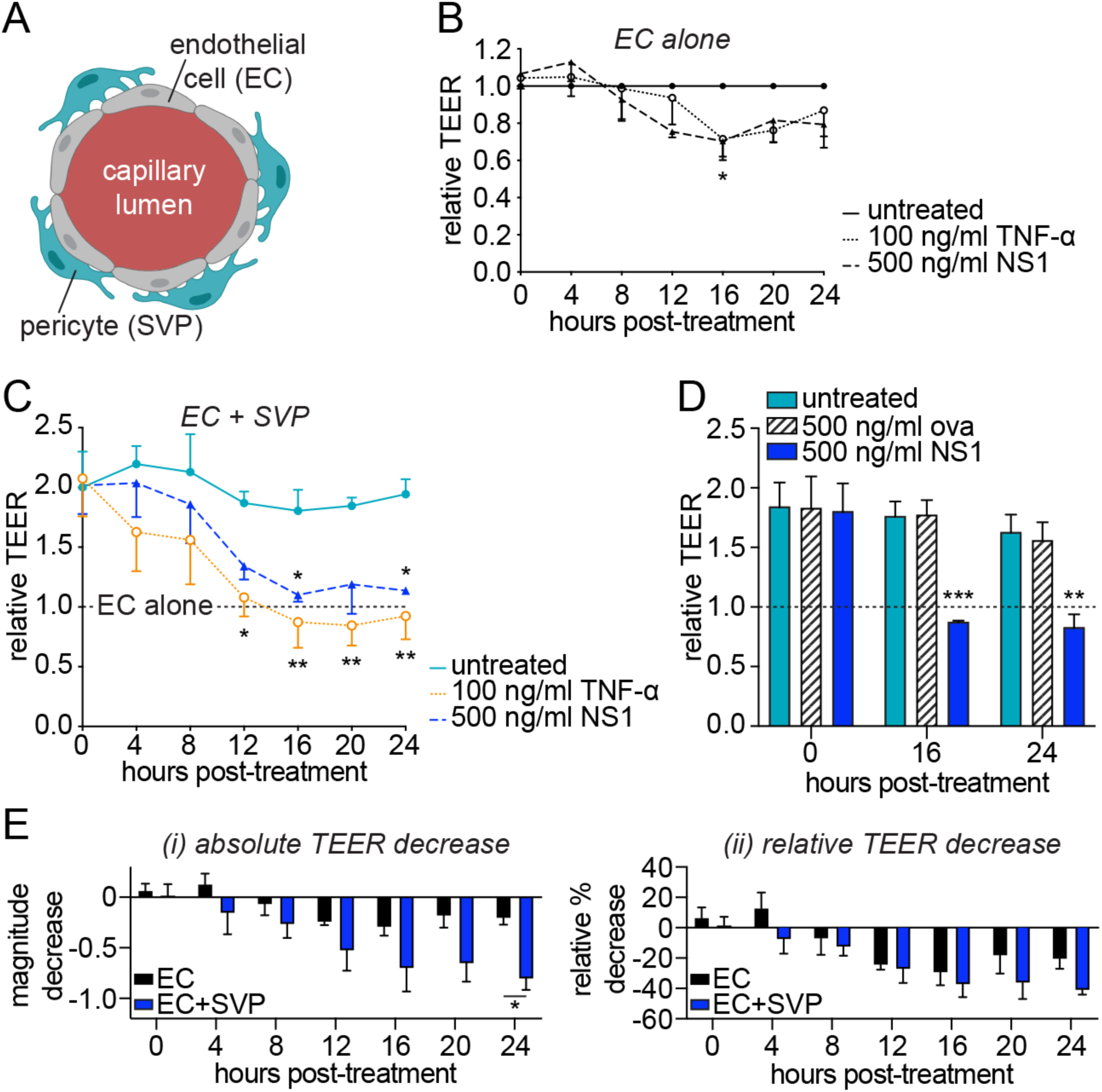
NS1 increases the permeability of an endothelial cell monolayer to a greater degree in the presence of co-cultured pericytes. (A) Cross-sectional illustration of the microvascular architecture, with pericytes (SVPs) supporting the maintenance of the endothelial barrier by endothelial cells (ECs). (B) Impact of treatment with purified recombinant TNF-α or DENV-2 NS1 on endothelial cell monolayer permeability over time as measured by TEER (trans-endothelial electrical resistance). At each individual time point, TEER values are normalised to untreated endothelial cells. (C and D) Enhancement of endothelial cell barrier function in co-culture with pericytes, and impact of treatment with DENV-2 NS1, TNF-α (C) or ovalbumin (“ova”) (D) on permeability of the co-culture over time. At each individual time point, TEER values are normalised to untreated endothelial cells cultured in the absence of pericytes (“EC alone”). (E) NS1-dependent decrease in TEER for endothelial cells cultured alone or in the presence of pericytes shown as absolute magnitude change in TEER (i) or percentage decrease relative to the respective untreated mono- or co-cultures (ii). All data minimum *N* = 3; n = 3. Error bars represent standard error of the mean. * *P* < 0.05; ** *P* < 0.01; *** *P* < 0.001. In Fig 1B, indicated significance is for NS1 treatment versus untreated endothelial cells. In Fig 1C and 1D, the difference between “EC alone” and “untreated” (co-cultures) is significant at *P* < 0.05. In Fig 1C, significant differences from “untreated” (co-cultures) are shown above the line for NS1 treatment and below the line for TNF-α treatment.

Here, we use microvascular hyperpermeability induced by DENV NS1 as a model to demonstrate a crucial role for pericytes in amplifying an infectious haemorrhagic syndrome. We show that DENV-2 NS1 induces hyperpermeability in *in vitro* co-cultures of primary pericytes and primary endothelial cells, and that the observed hyperpermeability is greater than for endothelial cells cultured alone. NS1 does not broadly affect all pericyte functions, but rather specifically reduces the capacity of pericytes to support endothelial cell function in 3D microvascular models. Finally, we demonstrate that NS1-induced hyperpermeability is not dependent on endothelial-pericyte cell contact and is at least partially mediated through effects on paracrine signalling. Importantly, the inclusion of pericytes in our *in vitro* model of dengue microvascular hyperpermeability reveals effects at markedly lower NS1 concentrations than in previously published studies, more representative of serum NS1 levels commonly observed in patients with severe dengue. Our findings could inform new strategies for developing diagnostics and treatments for severe dengue, and could have wide-reaching implications for the role of pericytes in other infectious vascular leakage syndromes.

## RESULTS

### NS1 triggers pericyte dysfunction and hyperpermeability in vascular cell co-cultures

We used trans-endothelial electrical resistance (TEER) as a measure of the strength of a cell monolayer barrier. In line with previous studies (11, 17), we found that treatment with DENV-2 NS1 reduced the barrier function of primary human umbilical vein endothelial cells (HUVECs) starting from 12 h post-treatment (Fig 1B). The magnitude of this NS1 effect was comparable to that of the vasodilatory cytokine TNF-α, here used as a positive control (Fig 1B).

Next, we wanted to explore whether pericytes, which are known to provide fundamental control of microvascular permeability by regulating endothelial cell function (22), contribute to NS1-induced hyperpermeability. Previously, we isolated a population of perivascular cells from the microvasculature surrounding human saphenous veins that expressed *bona fide* pericyte markers and supported endothelial cell function and angiogenesis *in vitro* and *in vivo* (26). Here, we established an *in vitro* model of the microvascular barrier in which HUVECs were co-cultured with saphenous vein pericytes (SVPs) in semi-contact, on either side of a porous membrane within a transwell system. Co-culture of endothelial cells with pericytes significantly increased endothelial barrier function compared to endothelial cells cultured alone, thus recapitulating the regulatory function of pericytes (Fig 1C). Treatment of these endothelial-pericyte co-cultures with 500 ng/ml DENV-2 NS1 significantly reduced endothelial barrier function (Fig 1C). This effect was evident from 8 h post-treatment and by 16 h the TEER of the co-cultures was indistinguishable from that of endothelial cells cultured alone. This indicates that NS1 completely abolished the ability of pericytes to support the barrier function of endothelial cells in this system. In contrast, ovalbumin did not affect permeability of the co-culture (Fig 1D), confirming the specificity of the observed pericyte dysfunction induced by DENV-2 NS1.

Notably, the absolute magnitude of the TEER decrease was significantly higher in the co-cultures compared to endothelial cells cultured alone, indicating a stronger effect of NS1 on permeability in the presence of pericytes (Fig 1Ei). Even when we accounted for the higher absolute TEER of the endothelial-pericyte co-cultures by normalising to the respective untreated controls, the decrease in barrier function in the co-culture was almost double that of endothelial cell monocultures at 20 h and 24 h post-treatment (Fig 1Eii).

Taken together, these results indicate that DENV-2 NS1 induces extensive dysfunction in primary human pericytes, leading to their inability to support endothelial cell barrier function. Furthermore, the effect NS1 has on permeability is more pronounced in the presence of pericytes, indicating that the presence of pericytes amplifies NS1-dependent microvascular hyperpermeability.

### NS1 disrupts the pericyte interaction with and support of the endothelial network

We next wanted to determine whether NS1 acts by disrupting the ability of pericytes to functionally support endothelial cells. Angiogenesis, the formation of new blood vessels from existing ones, is a complex process requiring precisely regulated reciprocal crosstalk between endothelial cells, which elongate and migrate to form new lumens, and pericytes, which ensheathe the branches and confer maturity to the new structures (27). Endothelial cells cultured in a 3D Matrigel substrate organise into capillary-like structures, and the maturation of these structures upon co-culture with pericyte is an established readout for the ability of pericytes to support endothelial cell and capillary functionality (26). In our hands, endothelial cells cultured on Matrigel formed abundant capillary-like structures after 6 h and the presence of pericytes visibly increased the thickness of the branches of these structures (Fig 2A). This increase in branch width is due to the physical association of pericytes with the endothelial structures, and the secretion of pro-angiogenic factors by pericytes directly stimulating the endothelium (26).

**Fig 2.**
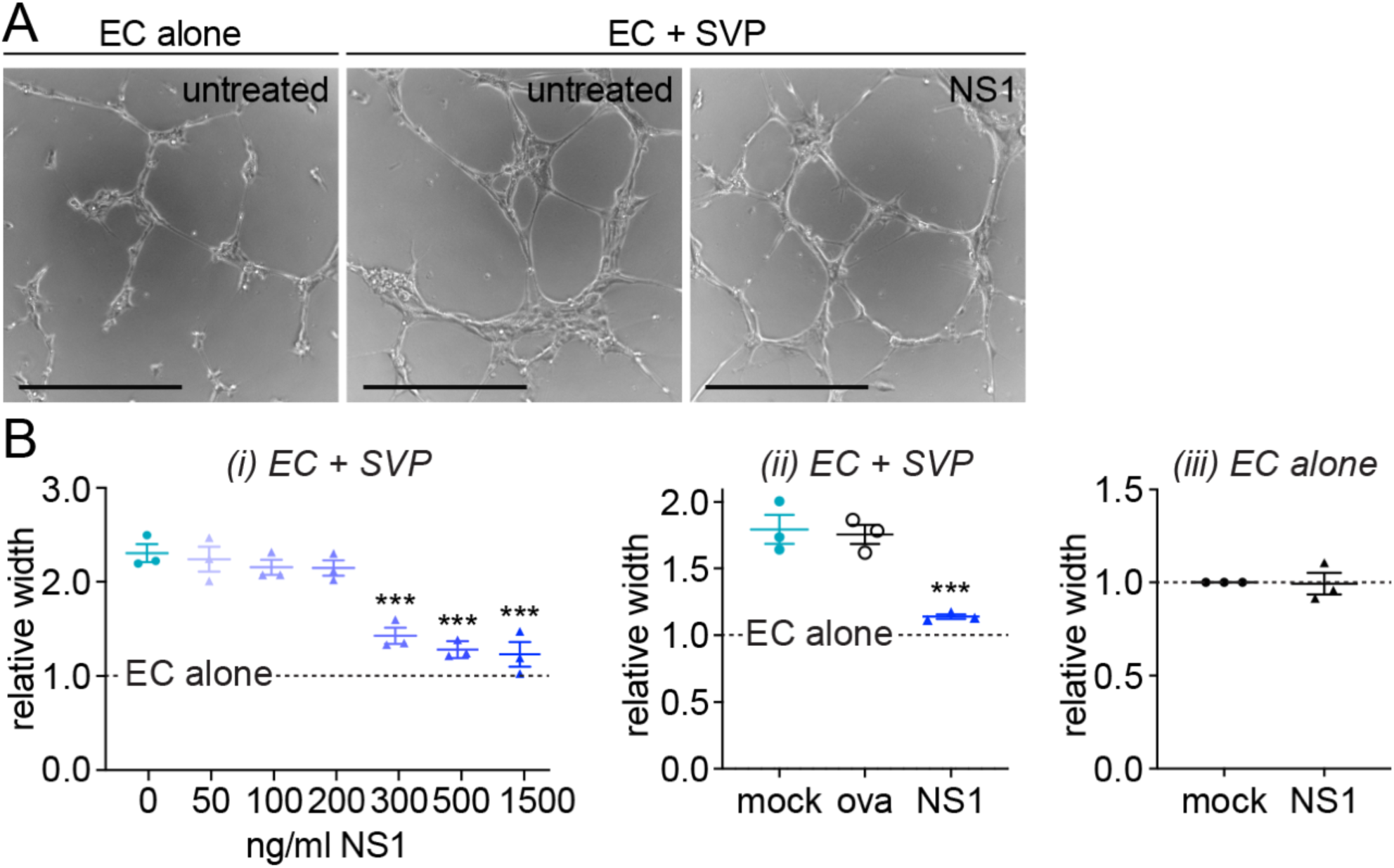
NS1 disrupts pericyte’s ability to support endothelial ‘capillary’ structures in a 3D microvascular model. (A) Representative images of endothelial cells forming capillary-like structures in 3D Matrigel culture in the presence (“EC + SVP”) or absence (“EC alone”) of pericytes, with or without treatment with 500 ng/ml DENV-2 NS1. Scale bars represent 400 µm. (B) Quantification of mean width of capillary-like structures in endothelial-pericyte co-cultures (i, ii) or endothelial cells cultured alone (iii) that were treated with NS1 or ovalbumin (“ova”) or left untreated (“mock”). Measurements are normalised to untreated endothelial cells cultured in the absence of pericytes (“EC alone”). Unless specified, recombinant protein concentration was 500 ng/ml. All data *N* = 3; n = 3. Error bars represent standard error of the mean. *** *P* < 0.001. In Fig 2Bi and 2Bii, significance shown is compared to untreated endothelial-pericyte co-cultures; the difference between “EC alone” and untreated co-cultures is significant at *P* < 0.0001.

Treatment of 3D endothelial-pericyte co-cultures with 500 ng/ml DENV-2 NS1 caused a reduction in branch width (Fig 2A), indicating a dysfunctional interaction between endothelial cells and pericytes. Quantification of NS1’s effect revealed a dose-dependent decrease in branch width, with doses above 300 ng/ml completely abolishing the contribution of pericytes to capillary-like structure formation (Fig 2Bi). In contrast, treatment with ovalbumin (“ova”) did not affect the endothelial-pericyte interaction in Matrigel (Fig 2Bii), demonstrating the specificity of the NS1 effect. Furthermore, branch width was not impacted by NS1 treatment when endothelial cells were cultured in the absence of pericytes, indicating that NS1 specifically interferes with the ability of pericytes to support the endothelium rather than the intrinsic angiogenic capacity of endothelial cells (Fig 2Biii). Taken together, our data suggest that the amplified effect NS1 has on endothelial permeability in the presence of pericytes is due to a disruption of pericyte functionalities required for maintenance of the microvascular architecture.

### NS1 does not broadly disrupt all pericyte functions

Our results demonstrate a robust effect of NS1 on pericyte function and in particular on the capacity of pericytes to support endothelial capillary-like structures and barrier function, two defining roles of pericytes. In order to exclude that the observed effect is due to a reduced viability of the pericyte or the endothelial populations, we treated each cell type separately with a range of NS1 concentrations (50-500 ng/ml). NS1 treatment for 48h did not affect cell growth or viability in either endothelial cells or pericytes (Fig 3A).

**Fig 3.**
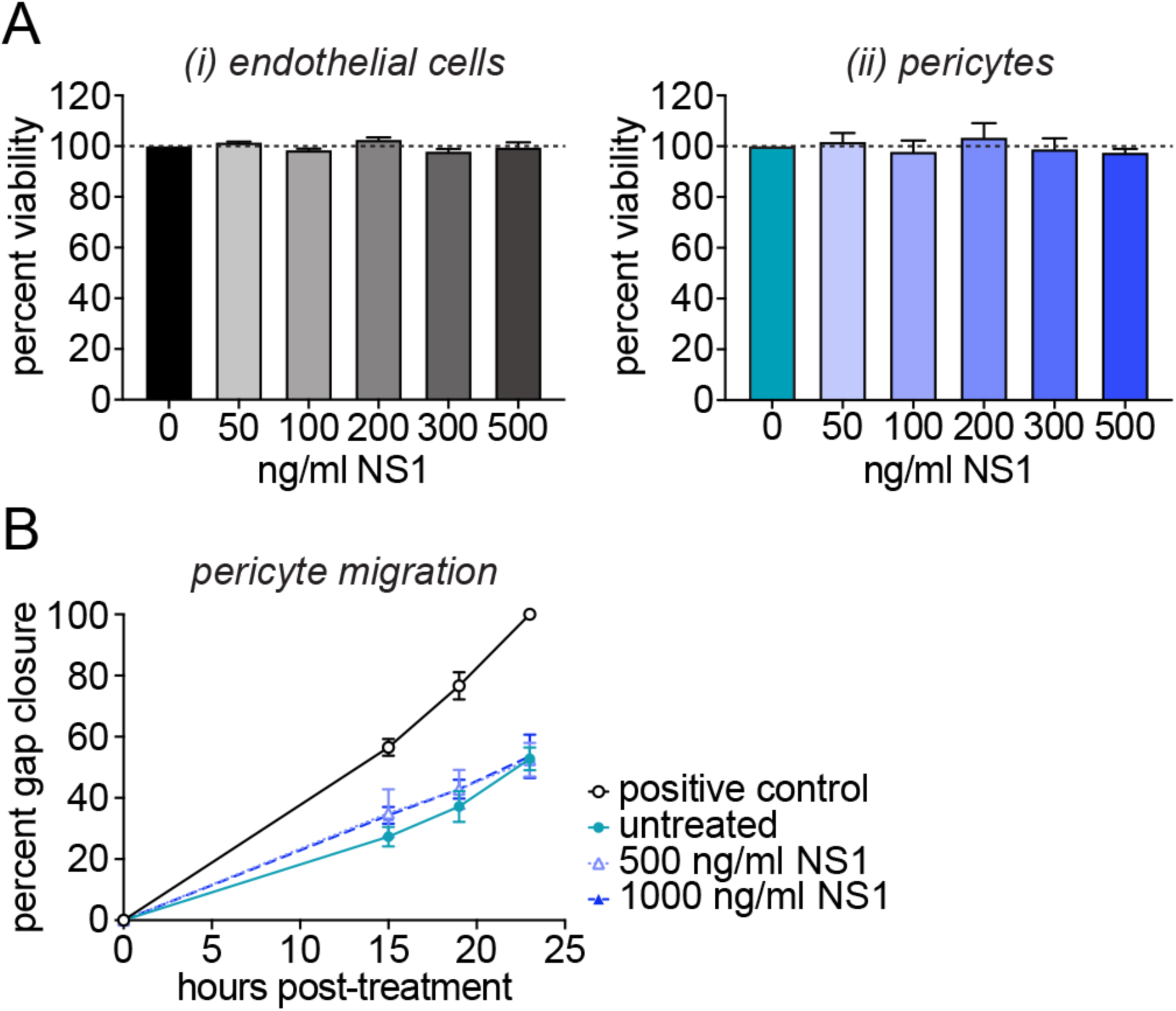
NS1 has no effect on pericyte viability or migration. (A) Impact of treatment with purified recombinant DENV-2 NS1 on the viability of endothelial cells (i) or pericytes (ii) 48 h post-treatment relative to untreated cells. *N* = 4, n = 6. (B) Impact of DENV-2 NS1 on the migration of pericytes. All values are normalised to the positive control 23 h post-scratch; positive control is pericytes grown in culture medium containing full complement of growth factors. *N* = 3, n = 3. Error bars represent standard error of the mean. Differences between treated and untreated cells are not statistically significant.

In order to support the microvascular architecture, pericytes must be able to migrate along capillaries. For this reason, we tested the intrinsic migratory capacity of pericytes upon treatment with NS1. DENV-2 NS1 concentration as high as 1,000 ng/ml had no effect on pericyte migration compared to the untreated control in a scratch assay measuring ‘wound’ closure following manual disruption of the cell monolayer (Fig 3B). These data further support our finding that NS1 specifically affects the endothelial-pericyte interaction.

### NS1 impacts pericyte paracrine signalling that supports the endothelial barrier

Pericytes regulate endothelial permeability through cell-cell interactions and paracrine signalling (22). In order to assess the paracrine contribution of pericytes to endothelial barrier function, we modified our TEER assay by seeding pericytes on the bottom of the well instead of on the membrane. This prevents any contact between endothelial cells and pericytes across the porous membrane. In this non-contact TEER, co-culture of endothelial cells with pericytes increased endothelial barrier function to a similar extent to that observed in semi-contact TEER (Fig 4A, compared to Fig 1C). This confirms that pericytes’ contribute to the capillary barrier is not simply as an additional physical layer, but as a master regulator of endothelial permeability. Treatment with 500 ng/ml DENV-2 NS1 decreased the resistance of the endothelial cell monolayer in the non-contact co-culture starting at 8 h post-treatment, and from 12 h post-treatment the contribution of pericytes to endothelial barrier function was completely ablated (Fig 4A). Furthermore, neither the absolute nor the relative effect size of NS1 treatment was significantly different when comparing the contact and non-contact co-cultures (Fig 4B), indicating that the effect of NS1 is mediated at least in part through paracrine modulators of endothelial barrier function.

**Fig 4.**
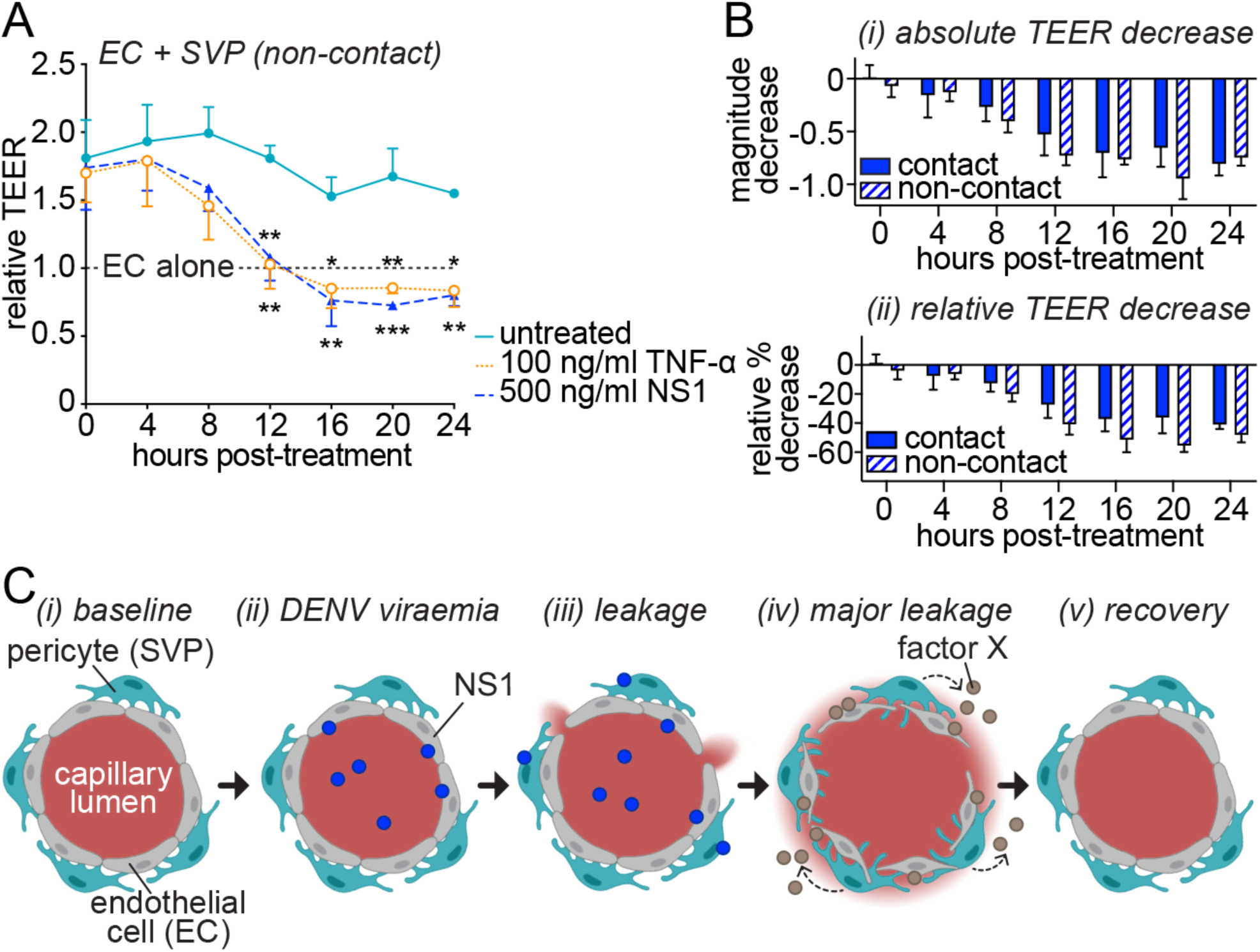
Pericytes amplify NS1-induced endothelial hyperpermeability via disrupted paracrine signalling. (A) Impact of treatment with purified recombinant TNF-α or DENV-2 NS1 on the permeability of endothelial cell monolayers co-cultured with pericytes without contact over time as measured by TEER (trans-endothelial electrical resistance). At each individual time point, TEER values are normalised to untreated endothelial cells cultured in the absence of pericytes (“EC alone”). (B) NS1-dependent decrease in TEER for endothelial-pericyte co-cultures grown with or without contact shown as absolute magnitude change in TEER (i) or percentage decrease relative to the respective untreated co-cultures (ii). (C) Working model for mechanism of pericyte amplification of NS1-induced endothelial permeability. (i) Under baseline conditions, pericytes support barrier maintenance by endothelial cells in the microvasculature. (ii) During DENV-2 infection, NS1 secreted into the circulatory system directly targets endothelial cells to (iii) induce vascular leakage, allowing NS1 to access pericytes as NS1 levels peak. (iv) Pericyte function is modulated by NS1, causing pericytes to secrete a paracrine factor (“factor X”) that signals to endothelial cells to further disrupt the endothelial barrier, resulting in enhanced vascular leakage after NS1 levels peak. (v) Pericytes recover from transient NS1 effects and normal endothelial-pericyte signalling is restored to allow cardiovascular recovery. All data *N* = 3; n = 4. Error bars represent standard error of the mean. * *P* < 0.05; ** *P* < 0.01; *** *P* < 0.001. In Fig 4A, the difference between “EC alone” and “untreated” (co-cultures) is significant at *P* < 0.05; significant differences from “untreated” (co-cultures) are shown above the line for TNF-α treatment and below the line for NS1 treatment.

## DISCUSSION

Here, we have described a role for pericytes in amplifying the hyperpermeability-inducing effects of DENV-2 NS1 on endothelial cells *in vitro*, and in doing so demonstrated that pericytes play a crucial role in the aetiology of an infectious haemorrhagic syndrome. NS1 had no effect on pericyte viability or migration, suggesting that NS1 specifically reduces the capacity of pericytes to support endothelial cell functions required for microvascular barrier formation and maintenance. We furthermore demonstrated that the effects of NS1 are at least partially the result of altered paracrine signalling between pericytes and endothelial cells. Since pericytes are an essential component of the microvascular architecture, we propose that pericytes play a crucial role in the aetiology of severe dengue. Interestingly, the distantly related flavivirus Japanese encephalitis virus (JEV) was recently shown to infect pericytes in the blood-brain barrier, where pericytes are also fundamental in maintaining the integrity of the barrier, allowing the virus to gain access to the brain to cause encephalitis (28). Pericytes may therefore play a wider role in the pathogenesis of flaviviruses that lead to diverse symptomatic outcomes.

Our data strongly suggest that NS1 interferes with the dynamic crosstalk between endothelial cells and pericytes. The specific factors mediating the NS1-dependent hyperpermeability remain unknown. Among several vasoactive molecules reported to be overrepresented in dengue patients, vascular endothelial growth factor (VEGF) has been shown to be upregulated twenty-fold compared to healthy controls (6). VEGF is abundantly (but not exclusively) secreted by pericytes to control endothelial cell angiogenesis and negatively regulates pericyte function (29, 30). Platelet-derived growth factor BB (PDGF-BB), a critical regulator of pericyte association with the endothelium, was also upregulated 20-60-fold in dengue patients, with a further increase at defervescence in patients with severe dengue (31). This timing coincides with the development of severe symptoms and is suggestive of an endothelial response to pericyte dysfunction. Finally, low levels of angiopoietin 1 (Ang1) and elevated levels of its antagonist angiopoietin 2 (Ang2) have been associated with dengue vascular leakage (5, 32), and the Ang1:Ang2 ratio has been proposed as a diagnostic marker for patients at risk of developing severe dengue (33). Ang1 is secreted by pericytes and platelets to increase endothelial barrier function (27), while Ang2, produced by endothelial cells and affecting pericyte coverage, is known to cause pericyte dropout in diabetic retinopathy (27, 34). It is tempting to speculate that one or more of these factors clinically associated with severe dengue might be indicative of a perturbation of pericyte function by NS1 *in vivo*.

We propose a mechanism whereby, early during DENV infection, NS1 secreted into patient serum induces well-described direct effects on endothelial cells causing local permeability that allow NS1 to gain access to pericytes on the apical side of the endothelium (Fig 4i-iii). Subsequent effects of NS1 on pericytes cause a dysregulation of paracrine signalling between pericytes and endothelial cells that leads to pronounced and potentially life-threatening hyperpermeability that manifests during defervescence (Fig 4iv). In patients that recover, the integrity of the microvascular endothelium is restored as pericytes regain the ability to support endothelial cell functions crucial to endothelial barrier formation and maintenance (Fig 4v).

Concentrations as high as 15,000 ng/ml of NS1 have been observed in the serum of dengue patients, however patients with severe dengue most commonly have serum NS1 levels ranging between 10-1,000 ng/ml (12-14). In our hands, NS1 affected the ability of pericytes to support endothelial cell function in 3D microvascular co-cultures at concentrations as low as 300 ng/ml (Fig 2B). To our knowledge, this is the lowest concentration at which microvascular effects of NS1 have been demonstrated *in vitro*. Furthermore, in endothelial-pericyte co-cultures, the observed reduction in TEER upon treatment with 500 ng/ml of NS1 is dramatically larger compared to endothelial cells cultured alone (Fig 1E). We therefore propose that modelling the essential role pericytes play in maintaining the microvascular endothelial barrier improves the ability of *in vitro* models of dengue hyperpermeability to recapitulate the effects of NS1 at more patient-relevant NS1 concentrations. Studying the effects of NS1 in our improved endothelial-pericyte co-culture system may identify novel molecular pathways and cellular mechanisms of dengue microvascular leakage that could facilitate the development of new diagnostics and treatments for severe dengue. Furthermore, our findings pave the way for future studies into the contribution that pericytes may make to the pathogenesis of other infectious diseases manifesting with symptoms of microvascular leakage.

## MATERIALS AND METHODS

### Cell culture

Human umbilical vein endothelial cells (HUVECs) were purchased from Lonza (Basel, Switzerland). Saphenous vein pericytes (SVPs) were isolated from patient-derived saphenous vein adventitial vasa vasorum by enzymatic digestion using CD34+CD31-selection as described in our previous publication (26). Both cell types were cultured in endothelial growth medium-2 (EGM-2; PromoCell, Heidelberg, Germany) and maintained at 37°C with 5% CO_2_. SVPs were grown on 1% fibronectin and 0.05% gelatin coated flasks (v/v). Cells were grown to 80% confluence and then passaged and used up to passage 8 for experiments.

### Recombinant proteins

DENV-2 NS1 was purchased from the Native Antigen Company (Kidlington, UK) and used between 50 and 1,500 ng/ml as indicated in the figures and figure legends. Ovalbumin (Invivogen, San Diego, USA) was used as negative control at 500 ng/ml and TNF-α (Peprotech, Rocky hill, USA) as a positive control at 100 ng/ml.

### Matrigel angiogenesis assay

Each well of a 96 well plate was coated with 30 μl of growth factor-reduced Matrigel (Corning, Corning, USA) on ice and allowed to solidify at 37°C for 30 minutes. Single cultures of 2 × 10^4^ cells/well of HUVECs or co-cultures of 1.5 × 10^4^ cells/well of HUVECs and 0.5 × 10^4^ of SVPs were seeded on top of the Matrigel. Five brightfield images were taken from each well 6 h after treatment.

### Trans-endothelial electrical resistance (TEER)

Millicell transwell inserts for 24-well plates (0.4 µm pores, Merk Millipore, Watford, UK) were coated with 1% fibronectin on both sides. Single cultures were established by seeding 1 × 10^5^ HUVECs per insert on one side of the transwell. For contact co-cultures, 2.5 × 10^4^ SVPs per insert were seeded on the other side of the transwell, 24 h later. For non-contact co-cultures, SVPs were alternatively seeded at the bottom of the 24-well plate. Cells were cultured to confluence and 50% of the medium was changed every 48 hours. TEER readings were taken using a Millicell ERS-2 Voltohmeter (Merk Millipore) immediately after treatment (0 h) and every 4 h for 24 h thereafter. Absolute TEER was calculated by subtracting the blank and multiplying by the well area; relative TEER was derived by dividing all values by the value of the HUVEC mono-cultures at each time point.

### Migration assay (wound healing)

SVPs were seeded in 48-well plates and allowed to grow to confluence, at which point the monolayer was scratched using a 10 µl pipette tip. The culture medium was changed to remove detached cells and treatments as indicated in endothelial basal medium (EBM, PromoCell) in the presence of 2 mM hydroxyurea to prevent cell proliferation. Pictures of the scratches were taken in the same positions every 4 h. Gap closure was calculated as the percentage of the distance travelled by the cell front divided by the initial size of the gap and normalised to the positive control within each replicate. The positive control was pericytes grown in culture medium containing a full complement of growth factors.

### Cell viability assay

HUVECs and SVPs were seeded in 96-well plates at 7 × 10^3^ cells/well and 2 × 10^3^ cells/well respectively. After overnight adhesion, cells were treated with recombinant proteins for 24 or 48 h. AlamarBlue™ (ThermoFisher Scientific, Waltham, MA USA) was then added 1:10 into the culture medium and fluorescence measured after 3 h using a spectrophotometer (excitation: 545nm, emission: 584nm). Relative viability was calculated in relation to the negative control.

### Microscopy and image analysis

Images were captured using either Eclipse Ts2 (Nikon, Tokyo, Japan) or EVOS Cell Imaging System (ThermoFisher) inverted microscopes. All images for a single analysis were collected using the same settings. For 3D microvascular cultures in Matrigel, branch width values were calculated manually using Fiji (ImageJ) image analysis software (National Institutes of Health, Bethesda, MA USA) by taking the average of the width of the thickest and thinnest section of all the branches in each image.

### Statistical analysis

Values are reported as mean±SEM. Experiments were repeated with a minimum of three independent biological replicates and three technical repeats within each replicate. Statistical analyses were performed with GraphPad Prism 8 software (GraphPad Software Inc, San Diego, CA USA). For Fig 1B, 1C, 1D, 3B and 4A, statistical analysis was by two-way repeated-measures ANOVA with Dunnett correction for multiple comparisons. Fig 1E and 4B were analysed by two-way repeated-measures ANOVA with Sidak correction for multiple comparisons. Fig 2Bi, 2Bii and 3A were analysed by one-way ANOVA with Dunnett correction for multiple comparisons. Fig 2Biii was analysed by unpaired Student’s *t* test.

### Figures

Graphs were plotted in GraphPad Prism 8 software. Illustrations were created with BioRender.com (Toronto, ON Canada). Figures were prepared in Adobe Illustrator (Adobe Systems, San Jose, CA USA). Images were cropped, annotated and modified to optimise brightness and contrast only.

## ACKNOWLEDGEMENTS

We wish to thank Ms. Kamelia Radeva and Ms. Diana Luna Buitrago for their contribution to the project. This work was supported by Medical Research Council grant MR/R010315/1 to KM and a University of Surrey Doctoral College and Department of Chemical and Process Engineering studentship to VM. DM was supported by National Centre for the Replacement, Refinement & Reduction of Animals in Research grants NC/R001006/1 and NC/T001216/1 to PC.

